# Face Familiarity Revealed by Fixational Eye Movements and Fixation-Related Potentials in Free Viewing

**DOI:** 10.1101/2022.04.28.489860

**Authors:** Oren Kadosh, Yoram Bonneh

**Affiliations:** School of Optometry and Vision Science, Faculty of Life Sciences, Bar-Ilan University, Ramat-Gan, Israel; The Leslie and Susan Gonda multidisciplinary Brain Research Center, Bar-Ilan University, Ramat-Gan, Israel

## Abstract

Event-related potentials (ERPs) and the oculomotor inhibition (OMI) in response to transient visual stimuli are known to be sensitive to different stimulus properties, including attention and expectation. In natural vision, transient stimulation of the visual cortex is generated primarily by saccades. Recent studies suggest that the core EEG components in free viewing, induced by saccades (fixation-related potentials, FRPs), are similar to the ERP components with flashed stimuli. We have recently found that the OMI in response to flashed stimuli is sensitive to face familiarity. Here, we investigated whether fixation-related-potentials (FRPs) and microsaccade inhibition (OMI) in free viewing are sensitive to face familiarity. Observers (N=15) freely watched a slideshow of seven unfamiliar and one familiar world leader’s facial images presented randomly for 4-seconds periods, with multiple images per identity. We measured the occipital fixation-related N1 relative to the P1 magnitude as well as the associated fixation-triggered OMI. We found that the average N1 was significantly smaller and the OMI was shorter for the familiar face, compared with any of the 7 unfamiliar faces. Moreover, the P1 was suppressed across saccades for the familiar but not for the unfamiliar faces. Overall, the results indicate that the occipital FRP and the OMI in free viewing are sensitive to face familiarity; this could perhaps be used as a novel physiological measure for studying hidden suppressed memories.

## Introduction

Studying vision in the natural settings of free viewing has become more common in many recent electrophysiological studies, demonstrating the advantages as well as the limitations of this method (Nikolaev, Meghanathan, & van Leeuwen, 2016). In general, these studies show results that are consistent with the event-related measurements; however, none of these studies probed face familiarity. Unlike traditional ERPs that used briefly flashed visual transients presented at the observer’s central visual field, in natural settings the scene is scanned over time via saccades following a peripheral preview.

### Free viewing

Accumulating evidence from recent free-viewing studies suggests that the brain’s response following a saccade, termed Fixation-related Potentials (FRP), exhibits electrophysiological components very similar to ERPs. For example, the saccade-related occipital lambda response reflects the same information processing as the classic VEP P1 (Kazai & Yagi, 2003). Recent studies that examined the face-selective activity at the lateral temporo-occipital electrodes, N170 (Bentin, Allison, Puce, Perez, & McCarthy, 1996) found a resembling increased negativity for faces in free viewing conditions (Auerbach-Asch, Bein, & Deouell, 2019; Buonocore, Dimigen, & Melcher, 2020). More classic findings were replicated in free viewing conditions, such as centro-parietal P300 elicited by target detection in a visual search (Hiebel et al., 2018) and the N400 priming effect in natural reading (Olaf Dimigen, Sommer, Hohlfeld, Jacobs, & Kliegl, 2011; Niefind & Dimigen, 2016). Combining EEG and eye-tracking measurements to study face familiarity enabled us to cross examine eye movement and electrophysiological changes over time, which are influenced by habituation and prior knowledge.

### Oculomotor Inhibition

Microsaccades are miniature saccades, with an average size of <0.5 dva, generated by neural activity in the superior colliculus (SC) (Hafed & Krauzlis, 2012). They occur during fixation, with a rate of one or two per second. Microsaccades are known to be inhibited momentarily (Oculomotor-Inhibition, OMI) (Bonneh, Adini, & Polat, 2016; White & Rolfs, 2016; Ziv & Bonneh, 2021) by stimulus presentation with a later release latency affected by the stimulus properties of attention and expectation. Whereas stimulus saliency, such as high contrast, is known to shorten the inhibition (Bonneh, Adini, & Polat, 2015), prolonged inhibition was found in response to deviants (Valsecchi, Betta, & Turatto, 2007). Although most studies used flashed stimuli, we have recently found similar inhibitory patterns in free viewing in response to stimulus saliency (Kadosh & Bonneh, 2022a).

### Face familiarity

Faces are considered complex stimuli that are processed holistically in posterior brain regions including the occipital face area (OFA), the fusiform face area (FFA), and the posterior superior temporal sulcus (STS), which form a core system for encoding the visual appearance of faces (Haxby, Hoffman, & Gobbini, 2000). This activity is modulated by top-down feedback from personal knowledge and emotional responses to alter the processing of familiar faces (Gobbini & Haxby, 2007). Face familiarity is of a major interest in research on concealed memories; the Concealed Information Test (CIT) is used for testing the familiarity of suspects to specific people and objects (Rosenfeld, 2020). Our recent study found prolonged OMI at fixation for a masked familiar face (Rosenzweig & Bonneh, 2019), which allowed us to reliably detect identity in a concealed information test (Rosenzweig & Bonneh, 2020). Next, we will review in more detail the currently known oculomotor and the ERP measures of familiarity.

### Face familiarity using oculomotor measures

Most of the experiments examining ocular measures in response to face familiarity used flashed stimuli. Rosenzweig & Bonneh (2019) measured the oculomotor inhibition of both microsaccades and blinks in response to masked novel and universally learned (Rosenzweig & Bonneh, 2019) or recently learned (Rosenzweig & Bonneh, 2020) familiar faces, and found prolonged OMI for the familiar faces in passive viewing. However, few studies have focused on different aspects of gaze fixations; one study claimed that the first two fixations are critical for revealing familiarity (Schwedes & Wentura, 2019), and other studies suggested that fewer fixations and longer fixation durations are the key for familiarity (Millen & Hancock, 2019; Peth, Kim, & Gamer, 2013). In contrast, another study reported more fixations and longer fixation durations for familiar faces with a CIT paradigm (Nahari, Lancry-Dayan, Ben-Shakhar, & Pertzov, 2019); however, when participants had to memorize faces, familiar faces initially attracted their gaze, but later triggered fewer fixations and shorter fixation durations (Lancry-Dayan, Nahari, Ben-Shakhar, & Pertzov, 2018). Overall, the eye movement behavior in relation to face familiarity depended on the instructions given by the researcher.

### Face familiarity using EEG Measurements

Previous electrophysiological familiarity studies measured the late event-related (ERP) response including the N250 (Bentin & Deouell, 2000; Gosling & Eimer, 2011) and the P300 contextual response. The face-sensitive N170 component, which reflects the structural encoding of faces prior to person identification (Huang et al., 2017; Sagiv & Bentin, 2001) was also tested; however, there were conflicting results. The majority of the studies did not find a familiarity effect (Anaki, Zion-Golumbic, & Bentin, 2007; Bentin & Deouell, 2000; Eimer, 2000; Gosling & Eimer, 2011; R. N. Henson et al., 2003); nevertheless, a few studies found an effect showing either larger N170 magnitudes for familiar faces (Barragan-Jason, Cauchoix, & Barbeau, 2015; Caharel, Courtay, Bernard, Lalonde, & Rebaï, 2005; Caharel, Fiori, Bernard, Lalonde, & Rebaï, 2006; Caharel et al., 2002; Keyes, Brady, Reilly, & Foxe, 2010) or smaller than for unfamiliar ones (Jemel, Pisani, Calabria, Crommelinck, & Bruyer, 2003; Marzi & Viggiano, 2007).

### Current study motivation and novelty

To date, little or no research has focused on earlier visual responses regarding face familiarity. However, a few studies examined the occipital P1 and N1 components using unfamiliar faces as stimuli, suggesting an early coarse processing of faces prior to face identification (Itier & Taylor, 2002; Nakashima et al., 2008). Here, we studied the fixation-triggered early posterior components that reflect feature and structure visual processing, and the timing of microsaccades during fixation, while participants freely viewed large images of both famous and unfamiliar faces presented for several seconds. We expected to find differences in the modulation over time, across successive saccades of the early occipital as well as the oculomotor responses by prior knowledge in the context of familiarity. Thus, the aim of this study was to investigate whether the FRP and OMI in free viewing are sensitive to face familiarity.

## Methods

### Participants

A total of sixteen observers were recruited for the experiment: 8 females and 8 males, aged 21-44. One participant was omitted from the data analysis due to the low quality of the data (more than 50% bad data). All participants had normal or corrected-to-normal vision and were naïve to the purpose of the study. The experiments were approved by the Bar-Ilan Internal Review Board (IRB) Ethics Committee. All participants gave written informed consent and all the experiments were conducted according to the IRB guidelines.

### Apparatus

The study combines eye tracking and electrophysiology recordings in free viewing synchronized by a split trigger to both systems. A wireless 8-channel headset with dry electrodes (Cognionics) was used for the EEG recordings and the Eyelink 1000 plus (SR Research) for eye tracking, both with a sampling rate of 500 Hz. A 100 Hz calibrated 24-in FHD LCD monitor (Eizo Foris fg2421), and the in-house developed integrative stimulus presentation and analysis tool (PSY) were developed by Y.S. Bonneh. Stimuli were displayed at a distance of 0.6 m. We used a 35 mm lens positioned 0.52 m from the participant’s stabilized head using a chin rest. The experiment was administered in dim light. All recordings were done binocularly, with analyses done on data from the left eye. A standard 9-point calibration was performed before each session.

### Saccade and Microsaccade RT calculation

For the saccade detection, we used an algorithm introduced by Engbert and Kliegl (Engbert & Kliegl, 2003), which is based on eye movement velocity. Microsaccades were detected as movements exceeding 8 *SD* of the mean velocity in 2D velocity space, as in (Rosenzweig & Bonneh, 2019; Yablonski, Polat, Bonneh, & Ben-Shachar, 2017). A velocity range of 8°/s-150°/s, an amplitude range of 0.08-1°, and a minimum duration of 9 *ms* were allowed for the microsaccades. We calculated the Warren Sarle’s bimodality coefficient (*BC*) of the saccade amplitude data, which is associated with the data skewness and kurtosis using a Matlab function by Hristo Zhivomirov, (2021). *BC* has a range of 0 to 1, where values greater than ~0.555 (the value for the uniform distribution) indicate bimodal or multimodal data distributions (Pfister, Schwarz, Janczyk, Dale, & Freeman, 2013). Eye tracking epochs were extracted, triggered by saccade (>1 *dva*) landing time in a range of −0.2 *sec* to 0.8 *sec* relative to the fixation onset with some overlap between epochs. This was taken into consideration when computing the microsaccade reaction time (msRT). The Microsaccade Reaction Time (msRT) was calculated for each epoch relative to the fixation onset in a predefined time window, as the latency of the first microsaccade in that window. The first fixation per trial was always ignored to avoid the flash effect on the OMI. The microsaccade RTs (msRT) were averaged across the epochs of each condition within observers and then averaged across observers, with error bars computed across observers on demeaned (within observer) data, with a correction factor (multiplied by √ (n/ (n-1). This method for computing the error bars allows a better representation of within-participant effects (Cousineau & Morey’s method (Morey, 2008); see also Bonneh et al. (Bonneh et al., 2015)). The saccade reaction time (sacRT) was calculated as the time interval between the current fixation onset and the next fixation onset, including only MS-free fixations.

### Fixation-related potentials - FRP

FRP epochs were created, as was done for the eye-tracking data, triggered by the saccade (>1 *dva*) landing time in a range of −0.1 *sec* to 0.3 *sec*, relative to the fixation onset to minimize overlapping data between epochs. Overlapping data points with a proximal saccade were excluded on both epochs. The first fixation per trial was always ignored to avoid the flash effect. We focused on the occipital electrodes O1, O2 and computed the positive and negative peaks in a predefined time range. The P1 peak was measured using a 50-150 *ms* time range, and the N1 was measured in a 100-200 *ms* window with no baseline correction. We then calculated the baseline-corrected peak-to-peak N1 relative to the P1 magnitude (N_1_’ = P_1_ – N_1_).

Peak extraction was optimized by setting an individual time range for each observer at around their average peak latency, within the predefined time range, from all the conditions combined. This was done to avoid using a long time range to overcome the latency differences across observers, which would increase the false peak discoveries. Extreme value artefacts were not allowed using a peak magnitude threshold exceeding ±50 *μVolts*. We excluded epochs with blinks or microsaccades that occurred at less than 200 *ms* after fixation onset to avoid peak corruption. The data were filtered using a 0.1 *Hz* high-pass and 30 *Hz* low-pass cutoffs. The EEG signal had a built-in reference using channels placed on both ears.

To ensure that we used a similar number of epochs per participant, we used an estimation of an average of 3 saccades per second to include only the first 12 epochs per trial (trial duration = 4 *seconds*), in the final analysis.

### Statistical assessment

Usually the statistical analysis of the variance (*One-way ANOVA*) and the Tukey multiple comparisons post-hoc tests were performed using Matlab. We first verified that the msRT distributions of different conditions come from normal distributions with equal variance. Another statistical method that was used is the *Linear Mix Model* (*LMM*) (Hohenstein, Matuschek, & Kliegl, 2017). The responses were fitted to a simple model of maximum likelihood with the *serial saccade number* or the *saccade size* used as the predictor variable, and the observer’s variability was set as the random effect. In addition, we computed Pearson’s linear correlation coefficient (*r^2^*) for the group averages of each plot.

### Stimuli and procedure

Observers freely watched a slideshow of seven unfamiliar and one familiar world leader’s facial images presented randomly for four-*second* periods, with multiple different images per person (see Figure 1). All images were chromatic with 600×800 minimum resolution, taken from the internet and were radially cropped around the head with a radius of 10 *dva*. The use of 5 different images per identity was done to reduce the effects of low image attributes and facial expressions from a specific image. A task was not used and the observers were just instructed to freely inspect the images.

**Figure 1.**
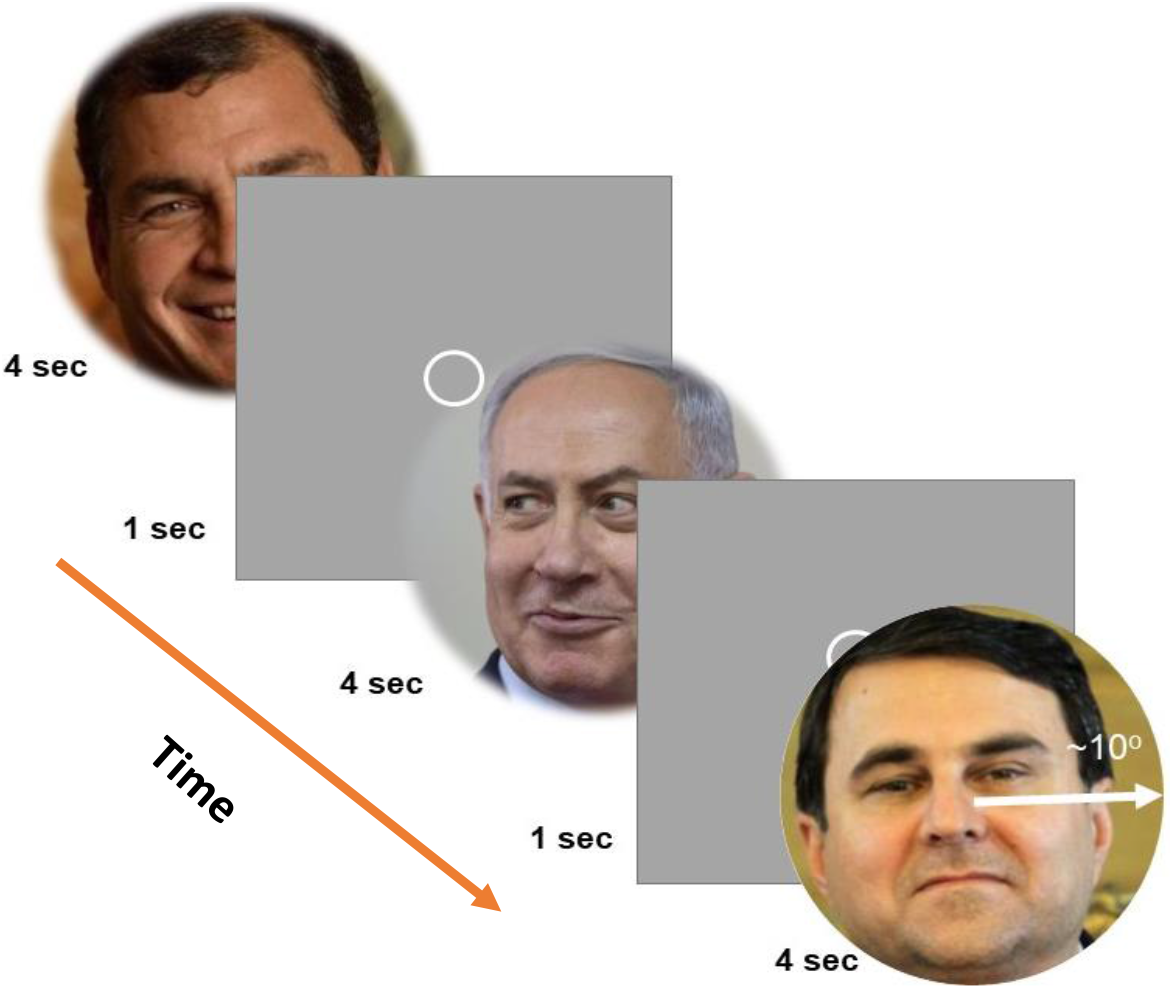
Stimuli and procedure. Observers (*N*=15) freely inspected large facial images (~10 *dva* radius, round images), presented in random order for four *seconds* each. In total, eight different identities were chosen of seven unfamiliar and one familiar world leader with five different images per identity.

## Results

We used static displays of faces, presented for four-second periods, while the observer viewed them freely without a task (see Figure 1, Stimuli and Procedure). Fixation-related potentials (FRPs), triggered by saccade landing time, were calculated, with a primary focus on early occipital components as well as Oculomotor-Inhibition (OMI), computed here as the timing of the first microsaccade following the fixation onset, excluding corrective microsaccades occurring proximate to the preceding saccade landing (see the Methods).

### EEG-FRP Results

The main familiarity effect was found at the posterior right hemisphere via the early occipital responses measured at the O2 electrode (see Figure 2a for a view of the EEG channel locations) with the peak magnitude computed (see the Methods) for the Lambda response (P1) peaking around ~90 *ms*, and the N1 peaking around ~140 *ms* after fixation onset (see Figure 2b, 2c). We then computed a baseline-corrected N1 by subtracting the P1 magnitude (peak-to-peak) per fixation-related epoch (see the Methods), which allowed us to test the combined FRP N1 and the P1 response (see Figure 3a). We hypothesized that the N1, which is often associated with a bottom-up prediction-error (PE), would reflect less PE for familiar faces with smaller magnitudes due to prior visual knowledge, which facilitates oculomotor dynamics and visual processing.

**Figure 2.**
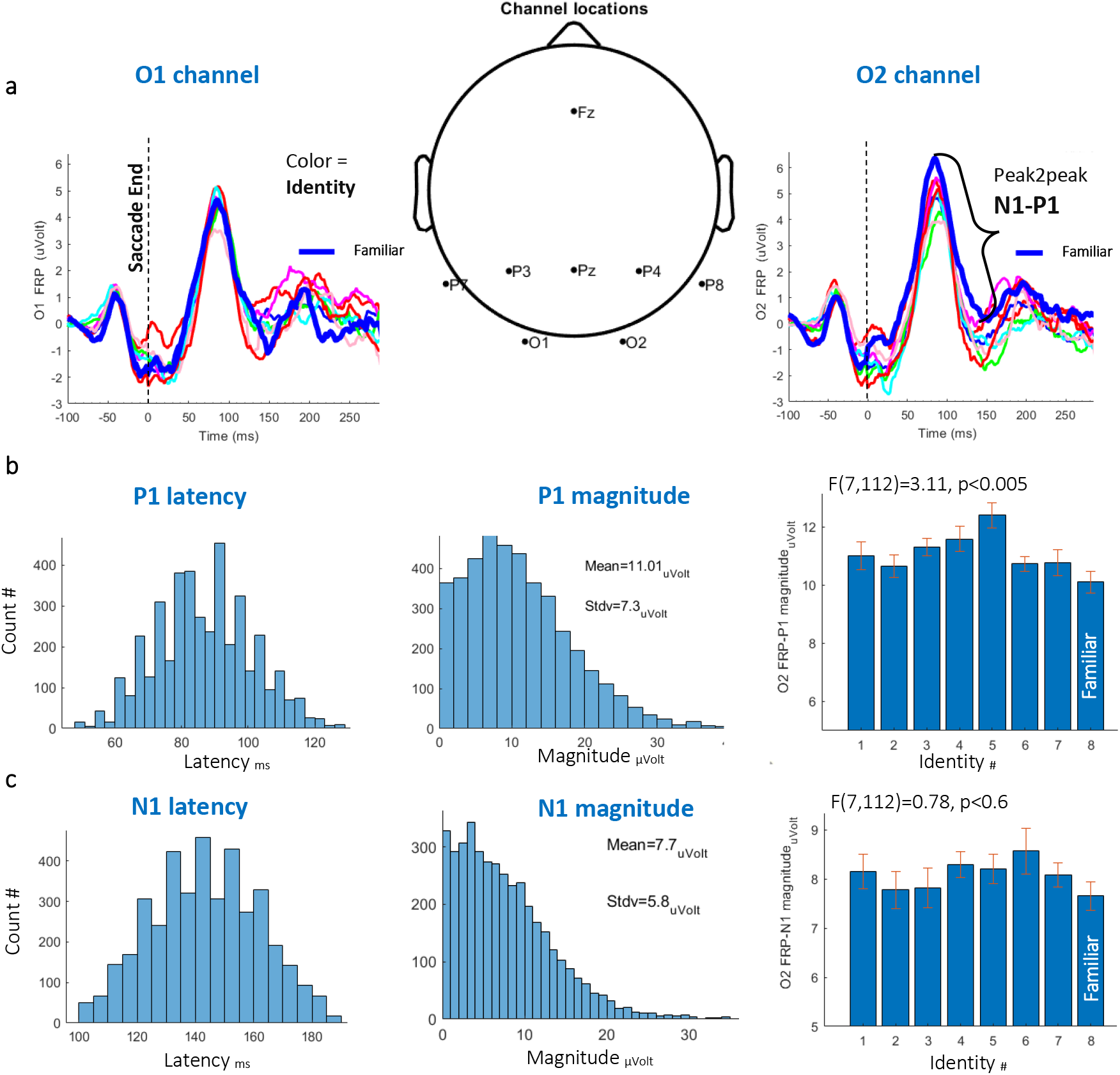
Basic FRP results. **a)** Scalp EEG channel topography and O1,O2 FRP results for all the unfamiliar faces (thin lines) and one familiar face (the thick blue line) averaged across observers (*N*=15). The vertical dashed line denotes the fixation onset. **b)** Results for the O2-FRP *P1 peak latency* (left), magnitude (center), and P1 averaged across observers’ magnitudes for each of the identities (right) with error bars calculated over the demeaned data showing a significantly smaller magnitude for the familiar identities (*F*(7,112)=3.11, *p*<0.005, *One-way ANOVA*). **c)** Results for O2-FRP *N1 peak latency* (left), magnitude (center), and N1 averaged across observers’ magnitudes for each of the identities (right) with error bars calculated over the demeaned data showing a non-significant smaller magnitude for the familiar identities (*p*<0.6).

**Figure 3.**
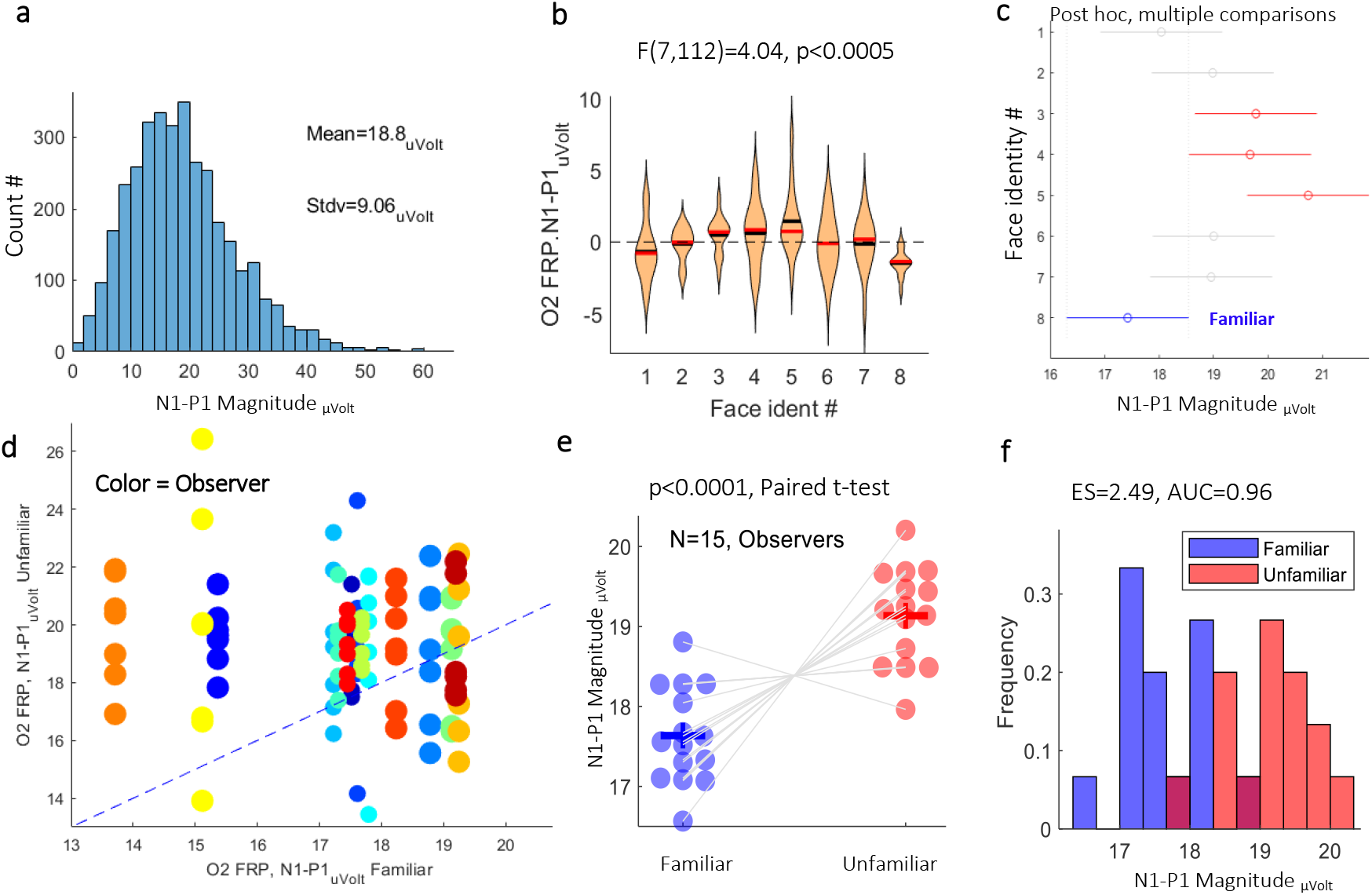
Normalized N1 familiarity effect. **a)** O2-FRP *N1 magnitude* histogram, all conditions combined (8 different facial identities). **b)** *N1 magnitude* for each of the 8 identities averaged across observers with error bars calculated over the demeaned data showing a significantly smaller magnitude for the familiar identity (*p*<0.0005, *One-way ANOVA*). **c)** Post-hoc multiple comparisons tests yielding three significantly different groups from the familiar identity, with 95%*confidence intervals*. **d)** A detailed observer scatter plot with a different color per participant and 7 dots of the unfamiliar identities; the *N1-magnitude* was compared with the familiar identity, showing that most of the dots are above the diagonal, signifying a larger magnitude for the unfamiliar identity. The dot size was reduced in crowded areas for better visibility. **e)** *Paired t-test* comparison between the familiar *N1 magnitude* and combined unfamiliar magnitudes showing a significant difference (*p*<0.0001). **f)** Histogram comparisons with the calculated effect size (*ES*=2.49, *Cohen’s d*) and the area under the ROC curve (*AUC*=0.96).

### The O2 FRP N1 familiarity effect

The results for *N1 magnitude* were calculated for all conditions (8 different face identities) averaged across observers (*N*=15) with error bars calculated over demeaned data. A histogram of the *bassline-corrected N1* is shown in Figure 3a. We found a significantly smaller *N1 magnitude* for the familiar identity, compared with each of the unfamiliar identities, *p*<0.0005 (*F*=(7,112)=4.04, *One-way ANOVA*, see Figure 3b). A multiple comparisons test yielded three out of seven significantly different groups from the familiar identity, with an illustration of the *confidence intervals* (see Figure 3c). To account for the individual contribution to the results, a detailed observer *scatter plot* with a different color for each participant and a dot for an unfamiliar identity *N1 magnitude*, compared with the familiar one, indicated that most of the dots are above the diagonal, signifying a larger magnitude for the unfamiliar one (see Figure 3d). To further demonstrate the individual results, we performed a *paired t-test* comparison between the familiar *N1 magnitude* and the combined unfamiliar magnitudes, which revealed a significant difference (*p*<0.0001, see Figure 3e). A histogram comparison, with a calculated huge effect size (*ES*=2.49, *Cohen’s d*) and an area under the ROC curve (*AUC*=0.96) is shown in Figure 3f.

### Adaptation effect on the FRP

We tested whether the adaptation of the occipital activity over successive saccades affects the FRP signal differently for the two categories. We found that the average *P1-magnitude* as a function of the *serial saccade number* was attenuated across successive saccades for the familiar identity via the O1 and O2 electrodes (*p*=0.04, *p*=0.0042, respectively. *Linear Mixed Model*), but not for the unfamiliar identity, with all identities combined (Figure 4a and Figure 4g). We then calculated the P1 (via O2) linear fit slopes separately for each of the identities and found that the average across observer negative slope was the largest for the familiar identity (Figure 4b). A histogram of the serial saccade number of the successive saccades per trial is shown in Figure 4c, with a mean of 7 ±3.6 *SD* saccades per trial. A detailed observer scatter plot with a different color for each participant and a dot for an unfamiliar identity slope value, compared with the familiar identity, showed that most of the dots are above the diagonal, indicating that the slope for the familiar identity was the most negative (see Figure 4d). A *paired t-test* comparison was performed between the familiar slope values and the combined unfamiliar ones, showing nearly a significant difference (*p*=0.062, see Figure 4e). A histogram comparison with a calculated medium effect size (*ES*=0.72, *Cohen’s d*) and the area under the *ROC* curve (*AUC*=0.66) is shown in Figure 4f. The *baseline-corrected N1* also shows a similar adaptation for the familiar identity (Figure 4h) as expected, because it was calculated relative to P1, peak-to-peak.

**Figure 4.**
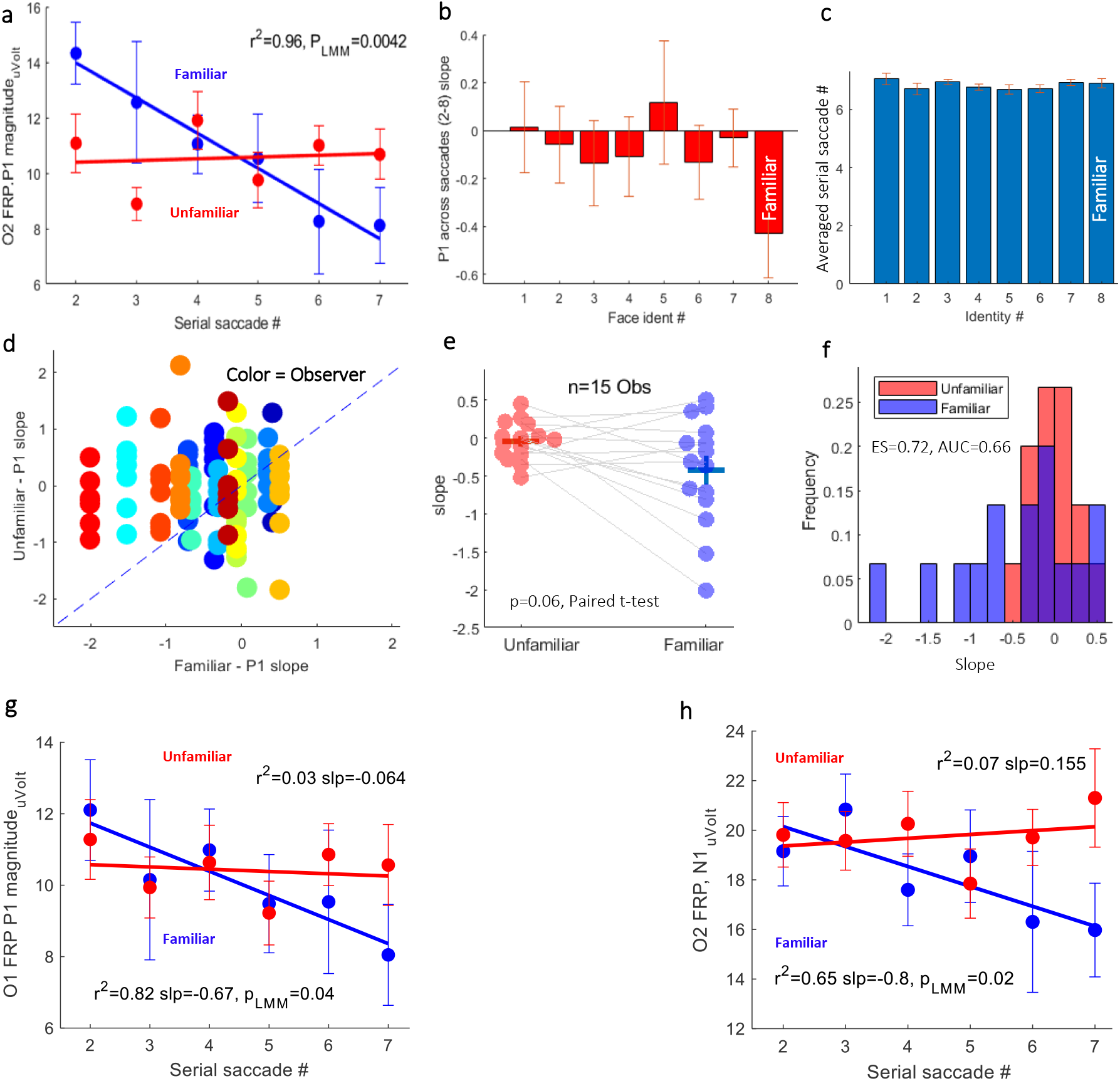
P1 adaptation effect. **a)** *O2 FRP P1 magnitude* as a function of the *serial saccade number* showing an adaptation effect across the first seven saccades, for the familiar (*p*=0.0042, *Linear Mixed Model*) but not for the unfamiliar identities combined. **b)** Slopes of the *O2 FRP P1 magnitude* linear fit across successive saccades for each of the identities. **c)** The *serial saccade number* of the successive saccades within a trial, averaged within and then across observers. **d)** Slopes of the *O2 FRP P1 magnitude* linear fit across successive saccades for each of the identities (dot) per observer (color); unfamiliar faces were compared with familiar identities, showing that most of the dots are above the diagonal, signifying a less negative slope for the unfamiliar identities. **e)** *Paired t-test* comparison between the familiar identities and the combined unfamiliar *P1 slopes* showing an almost significant difference (*p*=0.062). **f)** Histogram comparisons with the calculated medium *effect size* (*ES*=0.72, *Cohen’s d*) and the area under the ROC curve (*AUC*=0.66). **g)** *O1 FRP P1 magnitude* as a function of the *serial saccade number* showing an adaptation effect across the first seven saccades, for the familiar (*p*=0.04, *Linear Mixed Model*) but not for the unfamiliar identities combined. **h)** *O2 FRP N1 magnitude* as a function of the *serial saccade number* showing an adaptation effect across the first seven saccades, for the familiar (*p*=0.02, *Linear Mixed Model*) but not for the unfamiliar identities combined. In a-c, g-h, the data were averaged on observers and error bars were computed on the demeaned data.

### The Fixation-Related OMI familiarity effect

A recent study by Rosenzweig & Bonneh (2019) showed prolonged OMI for familiar faces briefly presented and masked. We wanted to determine whether microsaccade latencies after a saccade are also increased when a famous face is freely inspected for several seconds relative to an unfamiliar one. The results for the OMI are shown in Figure 5. Saccade and microsaccade size as well as the fixation duration histograms for all observers and conditions combined are shown in Figure 5a. The mean fixation duration was 350 *ms* ±175 *SD*. The saccade size distribution reflects a bimodal distribution of saccades and microsaccades with a calculation of the *bimodality coefficient* yielding a significance, *BC*=0.6 (see the Methods). The mean microsaccade size was 0.42 *dva* ±0.24 *SD*. We computed the msRT (microsaccade reaction time, see the Methods) as the first microsaccade (<0.6 *dva*) occurrence from 150 *ms* after the fixation onset, ignoring the corrective microsaccades that may occur immediately after the saccade. Figure 5b shows the average msRT across observers for each of the identities, showing decreased OMI for the familiar identity, which is opposite to the results obtained with flashed faces (Rosenzweig & Bonneh, 2019). Significance was assessed using *one-way ANOVA, F*(7,112)=2.84 and *p*<0.0093. The *post-hoc Tukey* method tests indicated that the familiar identity was significantly different only from one other unfamiliar group. The observer was denoted by color, in the *scatter plot* (Figure 5c), with a dot per identity, compared with the familiar identity. Interestingly, it was shown that most observers had longer msRTs for the unfamiliar identity (the dots above the symmetry line). Finally, the observer plot in Figure 5d compares the msRT in response to the familiar, relative to all unfamiliar identities combined and shows that 12 out of 15 observers had longer msRTs for the unfamiliar identities.

**Figure 5.**
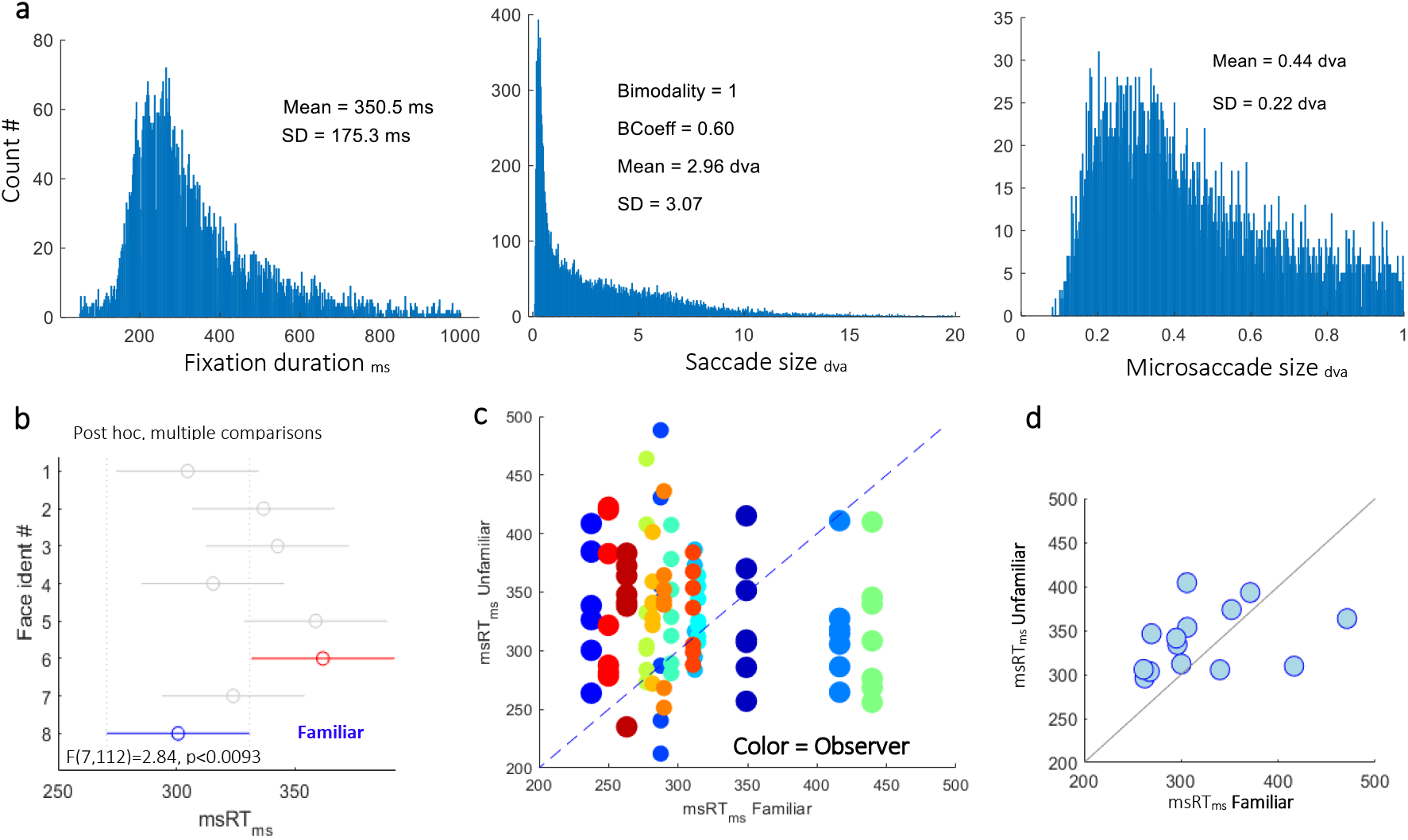
Oculomotor Inhibition (OMI) familiarity effect. **a)** Fixation duration, saccade, and microsaccades size histograms. **b)** OMI (msRT 150-800 *ms*) results for each of the 8 different face identities averaged across observers (*N=15*) showing a significantly shorter msRT for the familiar (*p*<0.0093, *One-way ANOVA, 95% confidence-intervals*). **c)** A detailed *scatter plot* with a different color per participant and a dot for an unfamiliar identity’s msRT compared with the familiar. Note that most of the dots are above the diagonal, signifying a longer OMI for the unfamiliar identity. The dot size was reduced in crowded areas for better visibility. **d)** Observer scatter plot comparing msRT for all unfamiliar identities combined vs. the familiar ones. As shown, 12 out of 15 observers were above the diagonal with longer msRT for the unfamiliar identities.

### The effect of saccade size

Previous FRP studies reported that the Lambda response (P1) amplitude increases with larger saccades (O. Dimigen, Valsecchi, Sommer, & Kliegl, 2009; Nikolaev et al., 2016; Ries, Slayback, & Touryan, 2018). This suggests that our finding of a decreased P1-N1 peak-to-peak magnitude for the familiar identity could result from smaller saccades that reduce the P1 amplitude. We therefore examined the effect of *saccade size* on the FRP. Figure 6 shows the effect of inducing the *saccade size* on the FRP and the relationship between the saccade size and familiarity. The saccade size histogram is shown in Figure 6a with a mean *saccade size* of 4.5 *dva* ±3 *SD*. As shown, the saccade size (>1 *dva*) did not differ between familiar and unfamiliar identities when averaged across observers (Figure 6b), or when plotted for each observer in a scatter plot (Figure 6c). A significant positive relation (*r^2^=0.41, Pearson correlation*) of the *P1 magnitude* and the *saccade size* (*p*=0.0016, *LMM*) is plotted in Figure 6d, which is consistent with previous studies. Figure 6e shows that the *N1-magnitude* was also positively correlated with saccade size (*r*^2^=0.36, *Pearson correlation;p*=0.012, *LMM*), because it was calculated relative to the *P1 magnitude* (peak-to-peak). Finally, the corrective microsaccade latencies show a negative correlation with *saccade size* (*r^2^*=0.85, *Pearson correlation; p*=0.00001, *LMM*); thus, larger saccades induced faster microsaccade reaction times due to the lower peripheral preview acuity (see Figure 6f).

**Figure 6.**
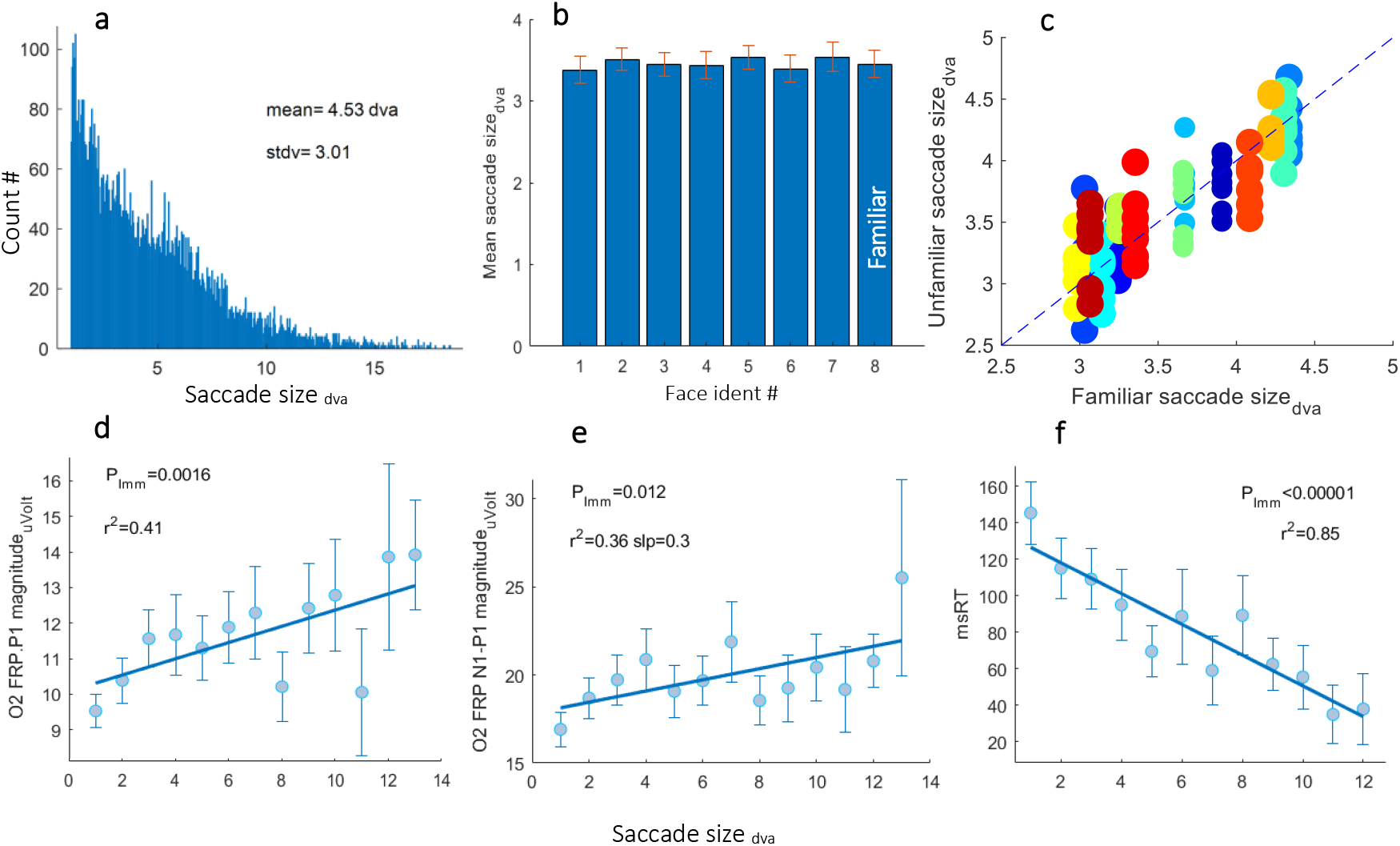
Saccade size effect. **a)** *Saccade size* (>1 *dva*) histogram for all observers and faces. **b)** Mean *saccade size* per identity, showing no difference between the familiar and unfamiliar identities. **c)** A detailed observer scatter plot comparing the average saccade size for the familiar identities with each of the unfamiliar ones, for each observer (a different color). As shown, *saccade size* did not differ with familiarity, indicated by the balanced distribution of the dots along the diagonal line. **d)** *P1-magnitudes* as a function of inducing the *saccade size*, averaged across observers, showing a significant positive relation (*p*=0.0016, *LMM*, see the Methods). **e)** FRP *N1 magnitude* was also positively correlated with *saccade size* (*r^2^*=0.36, *Pearson correlation; p*=0.012, *LMM*). **f)** Microsaccade latency (msRT) shows a negative correlation with the *saccade size* (*r^2^*=0.85, *Pearson correlation; p*=0.00001, *LMM*).

## Discussion

We investigated how familiarity in passive free viewing affected the early occipital responses that occur prior to face identification, and the effect on the latency of microsaccades, which is known to be affected by attention and expectations. Our results revealed a smaller fixation-related *N1 magnitude* and a shorter microsaccade inhibition (OMI) for the familiar faces, suggesting a lower prediction error while freely viewing familiar faces. Interestingly, the N1 FRP effect of familiarity appeared only at the right hemisphere, which is more specialized, according to some studies on face processing (Behrmann & Plaut, 2014; Yovel, Tambini, & Brandman, 2008).

We also examined the possible effect of adaptation and saccade size on the FRP and found that whereas saccade size did not differ between conditions and thus affected the FRP for different categories in the same way, the *P1 magnitude* was consistently adapted for the familiar faces and might have significantly contributed to the FRP familiarity effect. Next, we will discuss the three main factors that could affect the FRP results: (1) Prediction Error, PE, (2) Adaptation, and (3) Saccade size.

### With the help of prior knowledge: predictability/priming across saccades

We propose here that long-term familiarity provides a type of priming for the viewed image across saccades, revealed by a shorter OMI (see Figure 5) and smaller occipital P1 and N1 amplitudes (see Figure 3). These effects could be attributed to the effect of prior knowledge or long-term (over several seconds) priming by exposure to different photos having the same identity. Rare visual or auditory events are known to induce a longer OMI. For example, prolonged microsaccade inhibition has been reported for oddballs in a sequence, a rare blue patch among frequent red patches (Valsecchi et al., 2007), and for auditory deviants (Kadosh & Bonneh, 2022b; Valsecchi & Turatto, 2009; Widmann, Engbert, & Schroger, 2014). More preliminary evidence of prolonged inhibition was found for high-contrast patches among low-contrast patches (Bonneh et al., 2013) and for temporal oddballs via unpredicted intervals (Bonneh, Polat, & Adini, 2016). The ERP Mismatch-Negativity (MMN) is a well-known electrophysiological marker for oddball response. It reflects an automatic change detection mechanism in the auditory domain (Duncan-Johnson & Donchin, 1977; Giard, Perrin, Pernier, & Bouchet, 1990; Jääskeläinen et al., 2004; Näätänen, Gaillard, & Mäntysalo, 1978); it is manifested by a larger N1 negativity, peaking at about 170 *ms* at the temporal electrodes. Some studies suggest that a correlate of this prediction error generator also exists in the visual domain for visual mismatches termed vMMN (Stefanics, KremlÃ¡Ä• ek, & Czigler, 2014). Infrequent color patterns (Czigler, Balázs, & Pató, 2004), low spatial frequency gratings (Cleary, Donkers, Evans, & Belger, 2013), and face and house deviant orientations (Zhang et al., 2018) elicited a posterior vMMN. Even facial expressions and emotions were used in an oddball paradigm and elicited vMMN peaking at 100-200 *ms* (Astikainen, Cong, Ristaniemi, & Hietanen, 2013; Gayle, Gal, & Kieffaber, 2012; Kreegipuu et al., 2013).

All the above are examples of larger PE, measured by longer OMI and larger N1 amplitudes. Priming by repetition, on the other hand, has an effect that is opposite that of PE, manifested by attenuated responses similar to adaptation, and behavioral facilitation. Previous N170 ERP studies of face familiarity reported priming effects via smaller N170 amplitudes. For example, a study that used moony faces and priming by the same photo or a different photo, but with the same identity, found attenuated N170 amplitudes for the familiar identity, suggesting a top-down feedback effect, whereas a stronger priming effect was found with the same photo (Jemel et al., 2003). Short-term, within a second, priming effects, indicated by smaller N170 amplitudes, were reported in a face memory recall task for repeated stimuli, using inverted and contrast-reversed unknown faces (Itier & Taylor, 2002). Another study found identity-specific priming effects via attenuated P1 and N170 using morphed faces and argued that they may be due to low-level visual similarities (Walther, Schweinberger, Kaiser, & Kovács, 2013). A very recent study by Buonocore, Dimigen et al. found a reduction of the fixation-related N170, following an extra-foveal face preview, and contended that it is due to prediction (Buonocore et al., 2020).

In our recent paper, fixation-related OMI was measured for low-level stimuli in free viewing (Kadosh & Bonneh, 2022a). We found a resembling effect for OMI across saccades in the form of shortening of the OMI by repetition priming. Although this facilitation effect was found for gratings with the same spatial frequency, we believe that eye movement enhancement may occur due to familiarity or because different images have the same identity. Finally, we suggest that the N1 and OMI priming effects we found for the familiar identify did not occur independent of adaptation.

### Adaptation of the occipital P1 across saccades

Our results indicated that the occipital P1 for the familiar identity was attenuated across saccades (Figure 4). At least half of the participants showed a larger P1 adaptation for the familiar than for the unfamiliar identity. This attenuation could be interpreted as a simple reduction of attention for the familiar identity, because P1 reflects low-level features activity; this could also explain the shorter saccadic inhibition because attended stimuli induce longer OMI. However, it could also be related to prediction and priming. Adaptation and priming are difficult to separate; they can co-exist in the current settings and discriminating between the two requires further research. The process of attenuation over successive saccades is reminiscent of the repetition suppression phenomenon, which was found for face category and identity via fMRI and EEG. Image-invariant adaptation for familiar faces, but not for unfamiliar faces, was found in the face-selective regions of the medial temporal lobe, MTL, (Weibert et al., 2016). An fMRI study found a repetition suppression effect only for famous faces, whereas the opposite was found for unfamiliar ones (R. Henson, Shallice, & Dolan, 2000). More ERP studies found N170 adaptation to face category (Kloth, Schweinberger, & Kovács, 2010) as well as to face identity (Amihai, Deouell, & Bentin, 2011). A later identity-specific adaptation on the ERP was also found over superior occipito-temporal sites at around 200-280ms (Hills, Elward, & Lewis, 2010).

There is a known asymmetry of the ventral visual cortex for face processing in the literature. It is evident from lesions to the left/right hemispheres that affect word or face processing (Behrmann & Plaut, 2014; Yovel et al., 2008). In addition, the N170 also shows lateralization and is more prominent in the right hemisphere for faces than are words (Bentin et al., 1996; Maurer, Rossion, & McCandliss, 2008). In the current study, adaptation of the P1 FRP for the familiar identity appeared on both the O1 and O2 electrodes, placed at the posterior sites of the two hemispheres (see Figure 4a and 4g). However, the N1 FRP effect was found only through O2 (see Figure 3b). This can indicate that adaptation is not the main factor for the N1 FRP effect.

### Saccade size

In our study we found that the occipital *P1 magnitude* increased as a function of the amplitude of the saccade that precedes the fixation (see Figure 6) as previously reported (O. Dimigen et al., 2009; Nikolaev et al., 2016; Ries et al., 2018). This suggests that the effect of decreased *N1 magnitude* for the familiar identity could result from differences in the saccade amplitudes for different identities because the N1 negative peak value was baseline corrected by the preceding P1 peak value, peak-to-peak (see the Methods). For this reason, we measured the saccade size for each identity and found no significant difference between different identities (Figure 6b and 6c). The results also showed that the microsaccade timing after the saccade had a negative relationship with saccade size, meaning that after a large saccade, the corrective saccade was more immediate. In our OMI measurements we ignored the corrective saccades by including only microsaccade timings over 150 *ms* post-fixation onset.

### Comparison with previously concealed information from eye-movement studies

Previous familiarity studies that used eye tracking measurements detected fewer fixations with longer fixation durations when a familiar face was viewed (Nahari et al., 2019) and fewer areas of interest on the face when the participants were instructed to conceal their knowledge of familiarity (Millen & Hancock, 2019). With a memory task, familiar faces were less explored perhaps because they are easier to remember (Lancry-Dayan et al., 2018). In recent studies with different eye measurements, a longer OMI was found for microsaccades and for blinks when face stimuli were briefly presented and masked (Rosenzweig & Bonneh, 2019). Here we also used images with one familiar identity out of eight, but with a longer presentation duration. We expected to find similar results of longer OMI in free viewing, assuming that familiarity would raise associations, episodic-memories, and emotions, thus delaying the next saccade/microsaccade. However, we found the opposite from the expected, shorter microsaccade inhibition for the familiar identities, which might be related, as discussed before, to scanning enhancement due to priming/adaptation and feedback from other areas associated with prior knowledge.

### Conclusions

When observers free viewed a set of familiar and unfamiliar faces for a few seconds per face, the fixation-related potentials (FRPs) showed a decreased *N1 magnitude* for the familiar faces at the right occipital electrode and a shorter OMI for the familiar, compared with unfamiliar identities, both indicative of a smaller prediction error. The *P1 magnitude* for the familiar identity had been suppressed across successive saccades, implying priming or adaptation. In the current study, adaptation of the *P1 magnitude* for the familiar identity appeared on both hemispheres; however, the decreased *N1 magnitude* effect was observable only through the right occipital electrode, O2; this suggests that predictive structural face features, reflected by *N1* rather than adaptation of low-level features, reflected by *P1*, is the main factor driving this effect. Overall, the results indicate the sensitivity of the occipital FRP and the OMI in free viewing in relation to face familiarity; this could be used as a novel physiological measure for studying hidden memories.

## Author Contributions

OK and YSB designed the experiments. OK collected the data. OK and YSB developed the software used for running the experiments and the data analysis. OK analyzed the data and wrote the manuscript, YSB reviewed it.

## Data Availability

The experimental datasets generated during the current study will be available from the corresponding author upon reasonable request.

## References

Amihai, I., Deouell, L. Y., & Bentin, S. (2011). Neural adaptation is related to face repetition irrespective of identity: A reappraisal of the N170 effect. Experimental Brain Research, 209(2), 193–204. https://doi.org/10.1007/s00221-011-2546-x

Anaki, D., Zion-Golumbic, E., & Bentin, S. (2007). Electrophysiological neural mechanisms for detection, configural analysis and recognition of faces. NeuroImage, 37(4), 1407–1416. https://doi.org/10.1016/j.neuroimage.2007.05.054

Astikainen, P., Cong, F., Ristaniemi, T., & Hietanen, J. K.(2013). Event-related potentials to unattended changes in facial expressions: Detection of regularity violations or encoding of emotions? Frontiers in Human Neuroscience, 7(SEP), 1–10. https://doi.org/10.3389/fnhum.2013.00557

Auerbach-Asch, C. R., Bein, O., & Deouell, P. L. Y. (2019). Face Selective Neural Activity: comparison between fixed and free viewing.

Barragan-Jason, G., Cauchoix, M., & Barbeau, E. J. (2015). The neural speed of familiar face recognition. Neuropsychologia, 75, 390–401. https://doi.org/10.1016/j.neuropsychologia.2015.06.017

Behrmann, M., & Plaut, D. C. (2014). Bilateral hemispheric processing of words and faces: Evidence from word impairments in prosopagnosia and face impairments in pure alexia. Cerebral Cortex, 24(4), 1102–1118. https://doi.org/10.1093/cercor/bhs390

Bentin, S., Allison, T., Puce, A., Perez, E., & McCarthy, G. (1996). Electrophysiological studies of face perception in humans. Journal of Cognitive Neuroscience, 8(6), 551–565. https://doi.org/10.1162/jocn.1996.8.6.551

Bentin, S., & Deouell, L. Y. (2000). Structural encoding and identification in face processing: ERP evidence for separate mechanisms. Cognitive Neuropsychology, 17(1–3), 35–55. https://doi.org/10.1080/026432900380472

Bonneh, Y. S., Adini, Y., & Polat, U. (2015). Contrast sensitivity revealed by microsaccades. Journal of Vision, 15(9), 11. https://doi.org/10.1167/15.9.11

Bonneh, Y. S., Adini, Y., & Polat, U. (2016). Contrast sensitivity revealed by spontaneous eyeblinks: Evidence for a common mechanism of oculomotor inhibition. Journal of Vision, 16(7), 1.

Bonneh, Y. S., Adini, Y., Sagi, D., Tsodyks, M., Fried, M., & Arieli, A. (2013). Microsaccade latency uncovers stimulus predictability: Faster and longer inhibition for unpredicted stimuli. Journal of Vision, 13(9), 1342–1342. https://doi.org/10.1167/13.9.1342

Bonneh, Y. S., Polat, U., & Adini, Y. (2016). The buildup of temporal anticipation revealed by microsaccades and eye-blinks. Journal of Vision, 16(12), 935. https://doi.org/10.1167/16.12.935

Buonocore, A., Dimigen, O., & Melcher, D. (2020). Post-Saccadic Face Processing Is Modulated by Pre-Saccadic Preview: Evidence from Fixation-Related Potentials. The Journal of Neuroscience, 40(11), 2305–2313. https://doi.org/10.1523/JNEUROSCI.0861-19.2020

Caharel, S., Courtay, N., Bernard, C., Lalonde, R., & Rebaï, M. (2005). Familiarity and emotional expression influence an early stage of face processing: An electrophysiological study. Brain and Cognition, 59(1), 96–100. https://doi.org/10.1016/j.bandc.2005.05.005

Caharel, S., Fiori, N., Bernard, C., Lalonde, R., & Rebaï, M. (2006). The effects of inversion and eye displacements of familiar and unknown faces on early and late-stage ERPs. International Journal of Psychophysiology, 62(1), 141–151. https://doi.org/10.1016/j.ijpsycho.2006.03.002

Caharel, S., Poiroux, S., Bernard, C., Thibaut, F., Lalonde, R., & Rebai, M. (2002). ERPs associated with familiarity and degree of familiarity during face recognition. International Journal of Neuroscience, 112(12), 1499–1512. https://doi.org/10.1080/00207450290158368

Cleary, K. M., Donkers, F. C. L., Evans, A. M., & Belger, A. (2013). Investigating developmental changes in sensory processing: Visual mismatch response in healthy children. Frontiers in Human Neuroscience, 7(DEC), 1–13. https://doi.org/10.3389/fnhum.2013.00922

Czigler, I., Balázs, L., & Pató, L. G. (2004). Visual change detection: Event-related potentials are dependent on stimulus location in humans. Neuroscience Letters, 364(3), 149–153. https://doi.org/10.1016/j.neulet.2004.04.048

Dimigen, O., Valsecchi, M., Sommer, W., & Kliegl, R. (2009). Human Microsaccade-Related Visual Brain Responses. Journal of Neuroscience, 29(39), 12321–12331. https://doi.org/10.1523/JNEUROSCI.0911-09.2009

Dimigen, Olaf, Sommer, W., Hohlfeld, A., Jacobs, A. M., & Kliegl, R. (2011). Coregistration of eye movements and EEG in natural reading: Analyses and review. Journal of Experimental Psychology: General, 140(4), 552–572. https://doi.org/10.1037/a0023885

Duncan-Johnson, C. C., & Donchin, E. (1977). On quantifying surprise: the variation of event-related potentials with subjective probability. Psychophysiology, Vol. 14, pp. 456–467. https://doi.org/10.1111/j.1469-8986.1977.tb01312.x

Eimer, M. (2000). Event-related brain potentials distinguish processing stages involved in face perception and recognition. Clinical Neurophysiology, 111(4), 694–705. https://doi.org/10.1016/S1388-2457(99)00285-0

Engbert, R., & Kliegl, R. (2003). Microsaccades uncover the orientation of covert attention. Vision Research, 43(9), 1035–1045. https://doi.org/10.1016/S0042-6989(03)00084-1

Gayle, L. C., Gal, D. E., & Kieffaber, P. D. (2012). Measuring affective reactivity in individuals with autism spectrum personality traits using the visual mismatch negativity event-related brain potential. Frontiers in Human Neuroscience, 6(DEC), 1–7. https://doi.org/10.3389/fnhum.2012.00334

Giard, M.-H, Perrin, F., Pernier, J., & Bouchet, P. Brain Generators Implicated in the Processing of Auditory Stimulus Deviance: A Topographic Event-Related Potential Study., 27 Psychophysiology § (1990).

Gobbini, M. I., & Haxby, J. V. (2007). Neural systems for recognition of familiar faces. Neuropsychologia, 45(1), 32–41. https://doi.org/10.1016/j.neuropsychologia.2006.04.015

Gosling, A., & Eimer, M. (2011). An event-related brain potential study of explicit face recognition. Neuropsychologia, 49(9), 2736–2745. https://doi.org/10.1016/j.neuropsychologia.2011.05.025

Hafed, Z. M., & Krauzlis, R. J. (2012). Similarity of superior colliculus involvement in microsaccade and saccade generation. Journal of Neurophysiology, 107(7), 1904–1916. https://doi.org/10.1152/jn.01125.2011

Haxby, J. V, Hoffman, E. A., & Gobbini, M. I. (2000). The distributed human neural system for face perception. Trends Cogn Sci. 4:223-233, 4(6), 223–233.

Henson, R. N., Goshen-Gottstein, Y., Ganel, T., Otten, L. J., Quayle, A., & Rugg, M. D. (2003). Electrophysiological and haemodynamic correlates of face perception, recognition and priming. Cerebral Cortex, 13(7), 793–805. https://doi.org/10.1093/cercor/13.7.793

Henson, R., Shallice, T., & Dolan, R. (2000). Neuroimaging evidence for dissociable forms of repetition priming. Science, 287(5456), 1269–1272. https://doi.org/10.1126/science.287.5456.1269

Hiebel, H., Ischebeck, A., Brunner, C., Nikolaev, A. R., Höfler, M., & Körner, C. (2018). Target probability modulates fixation-related potentials in visual search. Biological Psychology, 138(May), 199–210. https://doi.org/10.1016/j.biopsycho.2018.09.007

Hills, P. J., Elward, R. L., & Lewis, M. B. (2010). Cross-modal face identity aftereffects and their relation to priming. Journal of Experimental Psychology: Human Perception and Performance, 36(4), 876–891. https://doi.org/10.1037/a0018731

Hohenstein, S., Matuschek, H., & Kliegl, R. (2017). Linked linear mixed models: A joint analysis of fixation locations and fixation durations in natural reading. Psychonomic Bulletin & Review, 24(3), 637–651. https://doi.org/10.3758/s13423-016-1138-y

Hristo Zhivomirov. (2021). Bimodality Coefficient Calculation with Matlab. Retrieved from https://www.mathworks.com/matlabcentral/fileexchange/84933-bimodality-coefficient-calculation-with-matlab

Huang, W., Wu, X., Hu, L., Wang, L., Ding, Y., & Qu, Z. (2017). Revisiting the earliest electrophysiological correlate of familiar face recognition. International Journal of Psychophysiology, 120(January), 42–53. https://doi.org/10.1016/j.ijpsycho.2017.07.001

Itier, R. J., & Taylor, M. J. (2002). Inversion and contrast polarity reversal affect both encoding and recognition processes of unfamiliar faces: A repetition study using ERPs. NeuroImage, 15(2), 353–372. https://doi.org/10.1006/nimg.2001.0982

Jääskeläinen, I. P., Ahveninen, J., Bonmassar, G., Dale, A. M., Ilmoniemi, R. J., Levänen, S.,… Belliveau, J. W. (2004). Human posterior auditory cortex gates novel sounds to consciousness. Proceedings of the National Academy of Sciences of the United States of America, 101(17), 6809–6814. https://doi.org/10.1073/pnas.0303760101

Jemel, B., Pisani, M., Calabria, M., Crommelinck, M., & Bruyer, R. (2003). Is the N170 for faces cognitively penetrable? Evidence from repetition priming of Mooney faces of familiar and unfamiliar persons. Cognitive Brain Research, 17(2), 431–446. https://doi.org/10.1016/S0926-6410(03)00145-9

Kadosh, O., & Bonneh, Y. S. (2022a). Fixation - related saccadic inhibition in free viewing in response to stimulus saliency. Scientific Reports, 1–12. https://doi.org/10.1038/s41598-022-10605-1

Kadosh, O., & Bonneh, Y. S. (2022b). Involuntary oculomotor inhibition markers of saliency and deviance in response to auditory sequences. Journal of Vision, 22(5), 8. https://doi.org/10.1167/jov.22.5.8

Kaunitz, L. N., Kamienkowski, J. E., Varatharajah, A., Sigman, M., Quiroga, R. Q., & Ison, M. J. (2014). Looking for a face in the crowd: Fixation-related potentials in an eye-movement visual search task. NeuroImage, 89, 297–305. https://doi.org/10.1016/j.neuroimage.2013.12.006

Kazai, K., & Yagi, A. (2003). the Lambda Response and the P100 Component. 3(1), 46–56.

Keyes, H., Brady, N., Reilly, R. B., & Foxe, J. J. (2010). My face or yours? Event-related potential correlates of self-face processing. Brain and Cognition, 72(2), 244–254. https://doi.org/10.1016/j.bandc.2009.09.006

Kloth, N., Schweinberger, S. R., & Kovács, G. (2010). Neural correlates of generic versus gender-specific face adaptation. Journal of Cognitive Neuroscience, 22(10), 2345–2356. https://doi.org/10.1162/jocn.2009.21329

Kreegipuu, K., Kuldkepp, N., Sibolt, O., Toom, M., Allik, J., & Näätänen, R. (2013). vMMN for schematic faces: Automatic detection of change in emotional expression. Frontiers in Human Neuroscience, 7(OCT), 1–11. https://doi.org/10.3389/fnhum.2013.00714

Lancry-Dayan, O. C., Nahari, T., Ben-Shakhar, G., & Pertzov, Y. (2018). Do You Know Him? Gaze Dynamics Toward Familiar Faces on a Concealed Information Test. Journal of Applied Research in Memory and Cognition, 7(2), 291–302. https://doi.org/10.1016/J.JARMAC.2018.01.011

Marzi, T., & Viggiano, M. P. (2007). Interplay between familiarity and orientation in face processing: An ERP study. International Journal of Psychophysiology, 65(3), 182–192. https://doi.org/10.1016/j.ijpsycho.2007.04.003

Maurer, U., Rossion, B., & McCandliss, B. D. (2008). Category specificity in early perception: Face and word N170 responses differ in both lateralization and habituation properties. Frontiers in Human Neuroscience, 2(DEC), 1–7. https://doi.org/10.3389/neuro.09.018.2008

Millen, A., & Hancock, P. (2019). Eye see through you! Eye tracking unmasks concealed face recognition despite countermeasures. https://doi.org/10.1186/s41235-019-0169-0

Morey, R. D. (2008). Confidence Intervals from Normalized Data: A correction to Cousineau (2005). Tutorials in Quantitative Methods for Psychology, 4(2), 61–64. https://doi.org/10.20982/tqmp.04.2.p061

Näätänen, R., Gaillard, A. W. K., & Mäntysalo, S. (1978). Early selective-attention effect on evoked potential reinterpreted. Acta Psychologica, 42(4), 313–329. https://doi.org/10.1016/0001-6918(78)90006-9

Nahari, T., Lancry-Dayan, O. C., Ben-Shakhar, G., & Pertzov, Y. (2019). Detecting concealed familiarity using eye movements: the role of task demands. Cognitive Research: Principles and Implications, 4(1). https://doi.org/10.1186/s41235-019-0162-7

Nakashima, T., Kaneko, K., Goto, Y., Abe, T., Mitsudo, T., Ogata, K.,… Tobimatsu, S. (2008). Early ERP components differentially extract facial features: Evidence for spatial frequency-and-contrast detectors. Neuroscience Research, 62(4), 225–235. https://doi.org/10.1016/j.neures.2008.08.009

Niefind, F., & Dimigen, O. (2016). Dissociating parafoveal preview benefit and parafovea-on-fovea effects during reading: A combined eye tracking and EEG study. Psychophysiology, 53(12). https://doi.org/10.1111/psyp.12765

Nikolaev, A. R., Meghanathan, R. N., & van Leeuwen, C. (2016). Combining EEG and eye movement recording in free viewing: Pitfalls and possibilities. Brain and Cognition, 107, 55–83. https://doi.org/10.1016/j.bandc.2016.06.004

Peth, J., Kim, J., & Gamer, M. (2013). Fixations and eye-blinks allow for detecting concealed crime related memories. https://doi.org/10.1016/j.ijpsycho.2013.03.003

Pfister, R., Schwarz, K. A., Janczyk, M., Dale, R., & Freeman, J. B. (2013). Good things peak in pairs: a note on the bimodality coefficient. Frontiers in Psychology, 4(1), 83–97. https://doi.org/10.3389/fpsyg.2013.00700

Ries, A. J., Slayback, D., & Touryan, J. (2018). The fixation-related lambda response: Effects of saccade magnitude, spatial frequency, and ocular artifact removal. International Journal of Psychophysiology, 134(August), 1–8. https://doi.org/10.1016/j.ijpsycho.2018.09.004

Rosenfeld, J. P. (2020). P300 in detecting concealed information and deception: A review. Psychophysiology, 57(7), 1–12. https://doi.org/10.1111/psyp.13362

Rosenzweig, G., & Bonneh, Y. S. (2019). Familiarity revealed by involuntary eye movements on the fringe of awareness. Scientific Reports, 9(1), 1–12. https://doi.org/10.1038/s41598-019-39889-6

Rosenzweig, G., & Bonneh, Y. S. (2020). Concealed information revealed by involuntary eye movements on the fringe of awareness in a mock terror experiment. Scientific Reports, 10(1), 1–15. https://doi.org/10.1038/s41598-020-71487-9

Sagiv, N., & Bentin, S. (2001). Structural encoding of human and schematic faces: Holistic and part-based processes. Journal of Cognitive Neuroscience, 13(7), 937–951. https://doi.org/10.1162/089892901753165854

Schwedes, C., & Wentura, D. (2019). The relevance of the first two eye fixations for recognition memory processes. https://doi.org/10.1080/09658211.2019.1567789

Stefanics, G., KremlÃ¡Ä•ek, J., & Czigler, I. (2014). Visual mismatch negativity: a predictive coding view. Frontiers in Human Neuroscience, 8(September), 1–19. https://doi.org/10.3389/fnhum.2014.00666

Valsecchi, M., Betta, E., & Turatto, M. (2007). Visual oddballs induce prolonged microsaccadic inhibition. Experimental Brain Research, 177(2), 196–208. https://doi.org/10.1007/s00221-006-0665-6

Valsecchi, M., & Turatto, M. (2009). Microsaccadic responses in a bimodal oddball task. Psychological Research, 73(1), 23–33. https://doi.org/10.1007/s00426-008-0142-x

Walther, C., Schweinberger, S. R., Kaiser, D., & Kovács, G. (2013). Neural correlates of priming and adaptation in familiar face perception. Cortex, 49(7), 1963–1977. https://doi.org/10.1016/j.cortex.2012.08.012

Weibert, K., Harris, R. J., Mitchell, A., Byrne, H., Young, A. W., & Andrews, T. J. (2016). An image-invariant neural response to familiar faces in the human medial temporal lobe. Cortex, 84, 34–42. https://doi.org/10.1016/j.cortex.2016.08.014

White, A. L., & Rolfs, M. (2016). Oculomotor inhibition covaries with conscious detection. Journal of Neurophysiology, 116(3), 1507–1521. https://doi.org/10.1152/jn.00268.2016

Widmann, A., Engbert, R., & Schroger, E. (2014). Microsaccadic Responses Indicate Fast Categorization of Sounds: A Novel Approach to Study Auditory Cognition. Journal of Neuroscience, 34(33), 11152–11158. https://doi.org/10.1523/JNEUROSCI.1568-14.2014

Yablonski, M., Polat, U., Bonneh, Y. S., & Ben-Shachar, M. (2017). Microsaccades are sensitive to word structure: A novel approach to study language processing. Scientific Reports, 7(1), 3999. https://doi.org/10.1038/s41598-017-04391-4

Yovel, G., Tambini, A., & Brandman, T. (2008). The asymmetry of the fusiform face area is a stable individual characteristic that underlies the left-visual-field superiority for faces. Neuropsychologia, 46(13), 3061–3068. https://doi.org/10.1016/j.neuropsychologia.2008.06.017

Zhang, Y., Li, J., Wang, Z., Zhao, L., Wang, W., & Miao, D. (2018). Perceptual expertise impacts preattentive processing of visual simple feature: A visual mismatch negativity study. NeuroReport, 29(5), 341–346. https://doi.org/10.1097/WNR.0000000000000947

Ziv, I., & Bonneh, Y. S. (2021). Oculomotor inhibition during smooth pursuit and its dependence on contrast sensitivity. Journal of Vision, 21(2), 1–20. https://doi.org/10.1167/jov.21.2.12

